# Astrocyte Regulation of Synaptic Plasticity Balances Robustness and Flexibility of Cell Assemblies

**DOI:** 10.1101/2024.08.08.607120

**Authors:** Roman Koshkin, Tomoki Fukai

## Abstract

Cell assemblies are believed to represent the substrate of memory. Although long-term plasticity likely enables the formation of cell assemblies, how other factors, such as astrocytes and short-term plasticity (STP), affect their properties is poorly understood. To close this gap, we investigated cell assembly dynamics in a recurrent network model mimicking the hippocampal area CA3. As shown in experiment, recurrent connections in our model obey a symmetric spike-timing-dependent plasticity (STDP), in which weight change may or may not depend on the releasable amount of neurotransmitter. The former case involves an interplay between STDP and STP. In addition, we implicitly modeled the effect of astrocyte NMDA receptors by manipulating the breadth of the distribution of neurotransmitter release probability in STP. Both STP-dependent and STP-independent STDP enabled spontaneous cell assembly formation. Under the former, however, cell assemblies tend to be smaller and more responsive to external stimulation, improving the network’s memory capacity and enabling flexible network restructuring. Furthermore, astrocyte regulation of the STP-dependent STDP facilitates stimulus-driven reorganization of neural networks without destroying existing assembly structure, thus balancing cell assemblies’ flexibility and robustness. Our findings elucidate the computational advantages of interaction between STP and STDP and highlight astrocytes’ possible regulatory role in memory formation.

## Introduction

A cell assembly – a strongly connected group of neurons – is believed to represent the fundamental building block of memory (Harris 2005; Josselyn and Tonegawa 2020). Models of cell assembly formation have shown that Hebbian plasticity is sufficient to explain stimulus-driven (Zenke, Agnes, and Gerstner 2015) and spontaneous (Triplett, Avitan, and Goodhill 2018) emergence of cell assemblies. However, despite much theoretical and in-vivo research, many questions remain about the conditions necessary for the emergence of cell assemblies and their properties. While some form of Hebbian plasticity appears to be sufficient for cell assembly formation, their properties – most interesting of which perhaps are stability and size – are shaped by multiple other factors and their interactions. For example, in some neurons the degree of weight change due to STDP was shown to depend on the availability of neurotransmitters (Froemke et al. 2006). This implies that there may exist an interaction between short-term plasticity (STP) and STDP. Further, *in vivo* experiments suggest that LTP is influenced by astrocytic NMDA receptors (Chipman et al. 2021), although the precise mechanism of this influence is not clear.

Here we investigate how cell assemblies emerge spontaneously, or driven by stimuli, under a symmetric STP-coupled STDP rule, in which the weight update depends not only on the relative timing of the post- and presynaptic spikes, but also on how much neurotransmitter is released by the presynapse. This rule was reported in the visual cortex (Froemke et al. 2006) and shown to facilitate sequence learning in a hippocampal network model (Haga and Fukai 2018). We also analyze how STDP kernel shape (determined by time constants) and release probability distribution affect the response of mature cell assembly structure to external stimuli. Finally, we implicitly modeled the effect of astrocytes on the network by manipulating the distribution of neurotransmitter release probability.

Astrocytes are a major glial cell type in the brain, and their interactions with neurons have attracted growing attention from researchers in experimental and computational neurosciences (Chen et al. 2023; Lyon and Allen 2022). A single astrocyte can interact with a vast number of synapses ranging from 100,000 in mice to 2 million in humans (Bushong et al. 2002; Oberheim et al. 2009) and regulate synaptogenesis (Baldwin and Eroglu 2017), synaptic pruning (Lee et al. 2021; Zhang et al. 2021), long-term synaptic plasticity and memory formation (Allen 2014; Allen and Eroglu 2017; Allen and Lyons 2018; Haydon and Nedergaard 2015; Lee and Chung 2019; Ota, Zanetti, and Hallock 2013; De Pittà, Brunel, and Volterra 2016; Santello, Toni, and Volterra 2019; Chung et al., 2015; Hösli et al., 2022; Sun et al., 2024; Akter et al., 2023; Sharma et al., 2023; Navarrete et al., 2019; Vignoli et al., 2021) and reward processing (Corkrum et al. 2020). Given the extensive scale of neuron-astrocyte interaction, astrocytes can substantially influence neural population dynamics by enhancing neuronal excitability (Bellot-Saez et al. 2017; Expósito 2024; Mizuno 2005), modulating correlated firing (Poskanzer and Yuste 2016; Delepine et al., 2023) and synaptic transmission (Deemyad, Lüthi, and Spruston 2018; Martin-Fernandez et al. 2017). Astrocytes are also implicated in neurodegenerative diseases and cognitive impairments (Richetin et al. 2020; Zimmer, Orr, and Orr 2024). One study revealed the crucial role of astrocytes in regulating short-term plasticity (STP) at cortical excitatory synapses. Namely, the blockade of NMDA receptors on astrocytes (aNMDAR) dramatically reduced the variance of neurotransmitter release probabilities across synapses, and simulations of a single neuron model suggested that widely varied release probabilities enhance Hebbian spike-timing-dependent plasticity of excitatory synapses (Chipman et al. 2021). STP significantly impacts spontaneous and stimulus-evoked dynamics of local cortical circuits, including their activity-dependent self-organization and reorganization processes. Understanding the interplay between astrocytic regulation and short-term plasticity is crucial for understanding astrocytes’ role in memory function.

Our simulations reveal that both STP-dependent and STP-independent symmetric STDP support the emergence of robust cell assemblies in a spiking neural network that receives no structured input. Interestingly, however, the STP-dependent STDP, but not STP-independent one, endows the spontaneous activity with two computationally desirable properties. First, it enables more stable cell assembly structure that better maintains self-similarity against representational drift (Driscoll et al. 2017; Mau, Hasselmo, and Cai 2020; Rule, O’Leary, and Harvey 2019) across time in the absence of structured stimulation. Second, the model becomes more responsive to external stimulation, making it easier for new stable cell assemblies to emerge. Finally, we found that the presence of astrocytes under STP-dependent STDP facilitates not only spontaneous self-organization but also stimulus-driven reorganization of the network without sacrificing the existing cell assembly structure. Our results underscore the computational advantage of interaction between STP and STDP and the crucial regulatory function astrocytes may play in memory formation.

## Results

The weights of excitatory recurrent connections in our network models undergo plastic changes obeying a symmetric STDP similar to that identified in the hippocampal CA3 (Mishra et al. 2016). While this hippocampal area is thought to engage in sequence generation (Buzsáki 2015; Davoudi and Foster 2019; Ecker et al. 2022), a symmetric STDP is unlikely to organize an asymmetric neuronal wiring supporting sequence generation. To reconcile this discrepancy between functional requirements and physiological observations, we examine two scenarios: one where the long-term plasticity rule (STDP) is coupled with short-term plasticity (STP), and another where it is not. The STP-coupled STDP, which was proposed previously to learn goal-directed sequential firing through reverse replay of hippocampal place cells (Haga and Fukai 2018), implies that the amplitudes of synaptic weight changes are proportional to neurotransmitter amounts available at the presynaptic release sites. In addition to the two STDP rules, we study the effect of astrocytes on the network dynamics by considering two additional cases: aNMDAR not blocked (astro+) and aNMDAR blocked (astro+). Thus, we compare the results of numerical simulations in the four cases arising from all possible combinations of (STP-independent and STP-dependent) × (astro- and astro+). In this study, the blockade of aNMDAR was indirectly modeled by adjusting the neurotransmitter release probability profile: broad in the astro+ condition and narrow in the astro-condition (refer to Materials and Methods for details).

### Spontaneous emergence of cell assemblies under the different STDP rules

Previous studies showed that STDP is sufficient for the emergence of cell assemblies (Hiratani and Fukai 2014). However, since the degree of weight change in our modified STP-dependent learning rule (Fig. 1B) depends on the amount of releasable neurotransmitter, we first verified that this modification does not interfere with the ability of our mode model (see Fig. 1A for the circuit diagram and Table 1 for detailed parameters) to form cell assemblies. Comparing the spike rasters at the beginning and end of the simulation (Fig. 1C) clearly indicates that the model’s spontaneous activity becomes organized under STP-dependent STDP. This is due to the formation of cell assemblies in the model’s weight matrix (Fig. 1D), which can be detected using a spectral clustering algorithm (refer to Materials and Methods for details).

**Figure 1.**
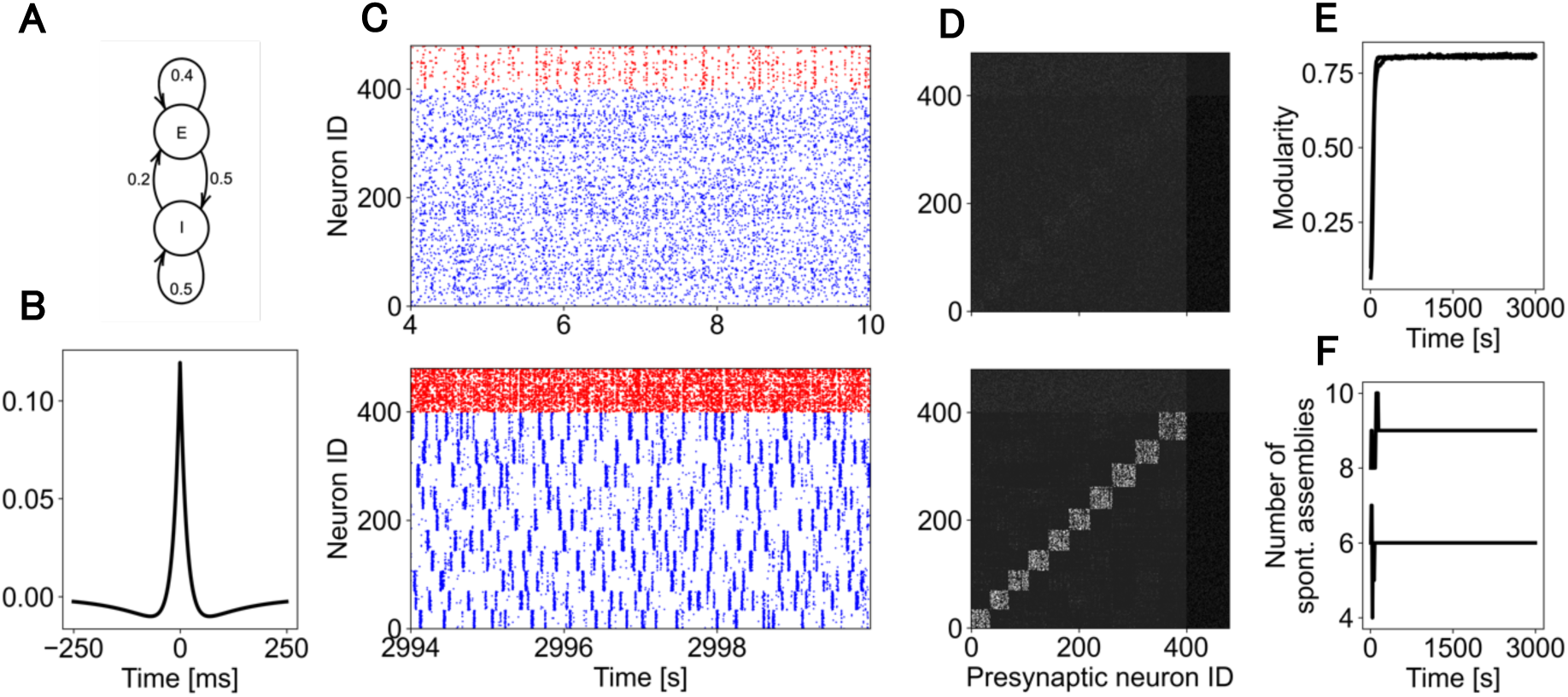
**Cell assemblies self-organized in the model with various parameter values**. The circuit diagram (A) and symmetric E-to-E STDP kernel (B) used in the simulation are shown. (C) Spike rasters show 6 seconds of spontaneous activity at the beginning (top) and end (bottom) of the simulation. Inhibitory neurons are shown in red, while excitatory ones in blue. In the bottom, excitatory neurons are rearranged to reveal a cell assembly structure. (D) The weight matrix at 10 and 2930 s of simulation (top and bottom, respectively). (E) The weight matrix modularity eventually plateaus, and (F) the number of cell assemblies also stabilizes. In E and F, 25 model simulations with different initial conditions are shown.

**Table 1.**
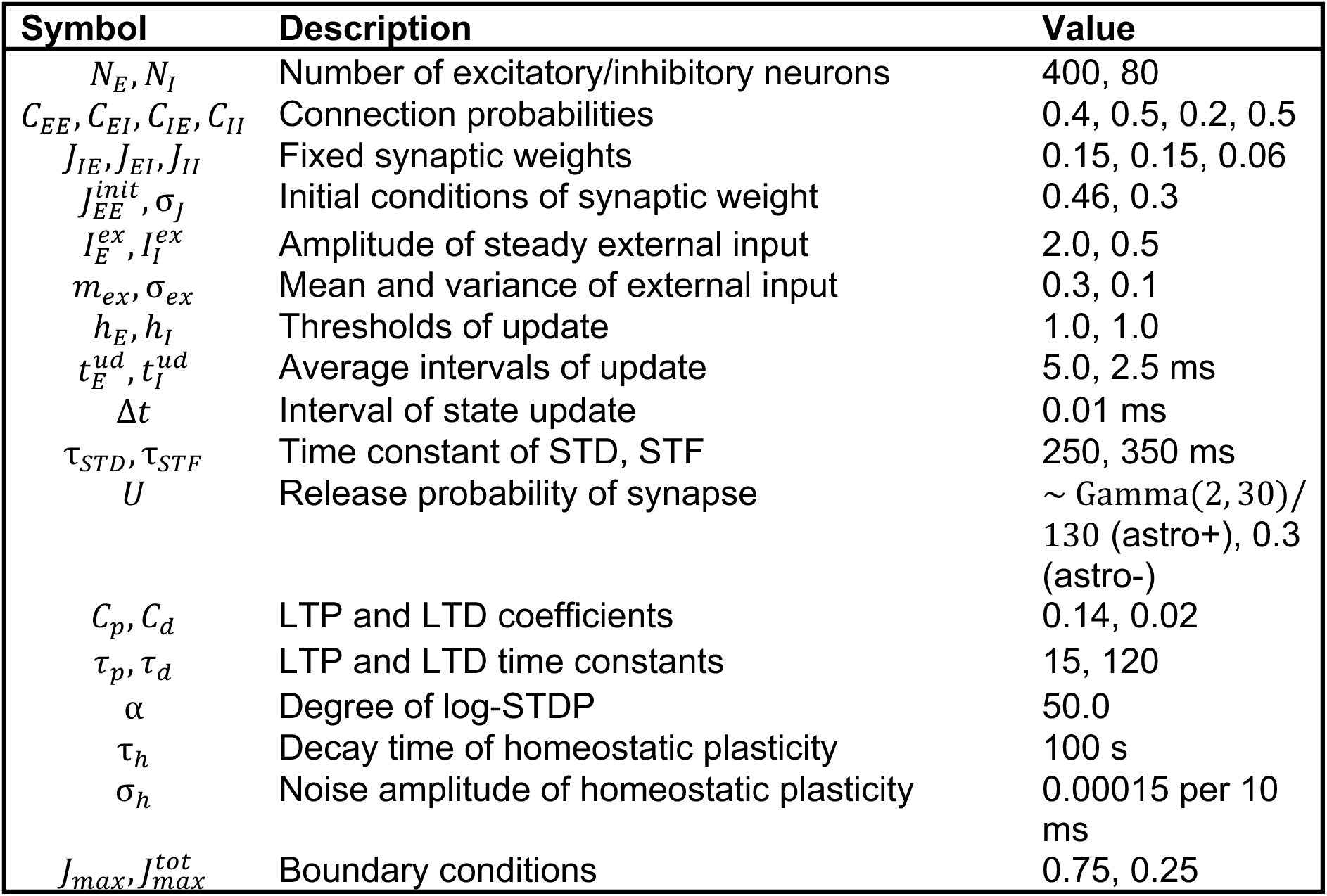
Model parameters.

We can quantify the degree of distinction between cell assemblies by the modularity of the connection matrix, which takes values between 0 and 1. The higher the modularity, the more distinct the individual cell assemblies are. Initially, the weight matrix has a low modularity (Fig. 1E) and contains spurious assemblies (Fig. 1F). As the modularity of the network grows to a sufficient value (about 0.2), the number of cell assemblies begins to stabilize and eventually reaches an equilibrium.

Both under STP-independent and STP-coupled STDP, the number of self-organized cell assemblies at equilibrium depended mostly on the coefficient of long-term depression (LTD) but not long-term potentiation (LTP), or short-term plasticity time constants (STF and STD). As expected, a network evolving under STP-independent STDP can also converge to a stable number of cell assemblies (Fig. 2). To see the influence of various parameters of learning rules, especially STP-dependence, on spontaneous formation of cell-assembles, we calculated the number of cell assemblies at equilibrium, i.e., when the number of spontaneously organized cell assemblies stabilized. To this end, we ran a grid of 1024 simulations of the same model, each with different values *C_d_* ∈ 0.0,0.3,0.5,0.7 and *C_p_* ∈ 0.05,0.1,0.15,0.2 (Fig. 2A and 2B, respectively) and STF (*τ_STF_* ∈ 100,200,300,400) and STD (*τ_STD_* ∈ 100,200,300,400) time constants (Fig. 2C and D, respectively). The models converged to different numbers of cell assemblies, despite having the same number of neurons and other parameters except *τ_STD_*, *τ_STF_*, *C_d_*and *C_p_*in each simulation. The modularity of the model was distributed broadly, revealing no simple relationships with the number of cell assemblies (Fig. 2E). Intriguingly, we found that the coefficient of LTD (*C_d_*), which changes the depth of the LTD component, is the strongest predictor of the number of spontaneously organized cell assemblies (Fig. 2A, F). However, the coefficient of LTP (*C_p_*), which changes the height of the LTP component, does not seem to clearly correlate with the number of cell assemblies (Fig. 2B, F). We speculate that this happens because LTD suppresses the average number of co-active neurons in a cell assembly, as implied in Fig. 1B (i.e., the number of assemblies and the average number of co-active neurons should be negatively correlated). We discuss this point in more detail in the section on cell assembly stability analysis.

**Figure 2.**
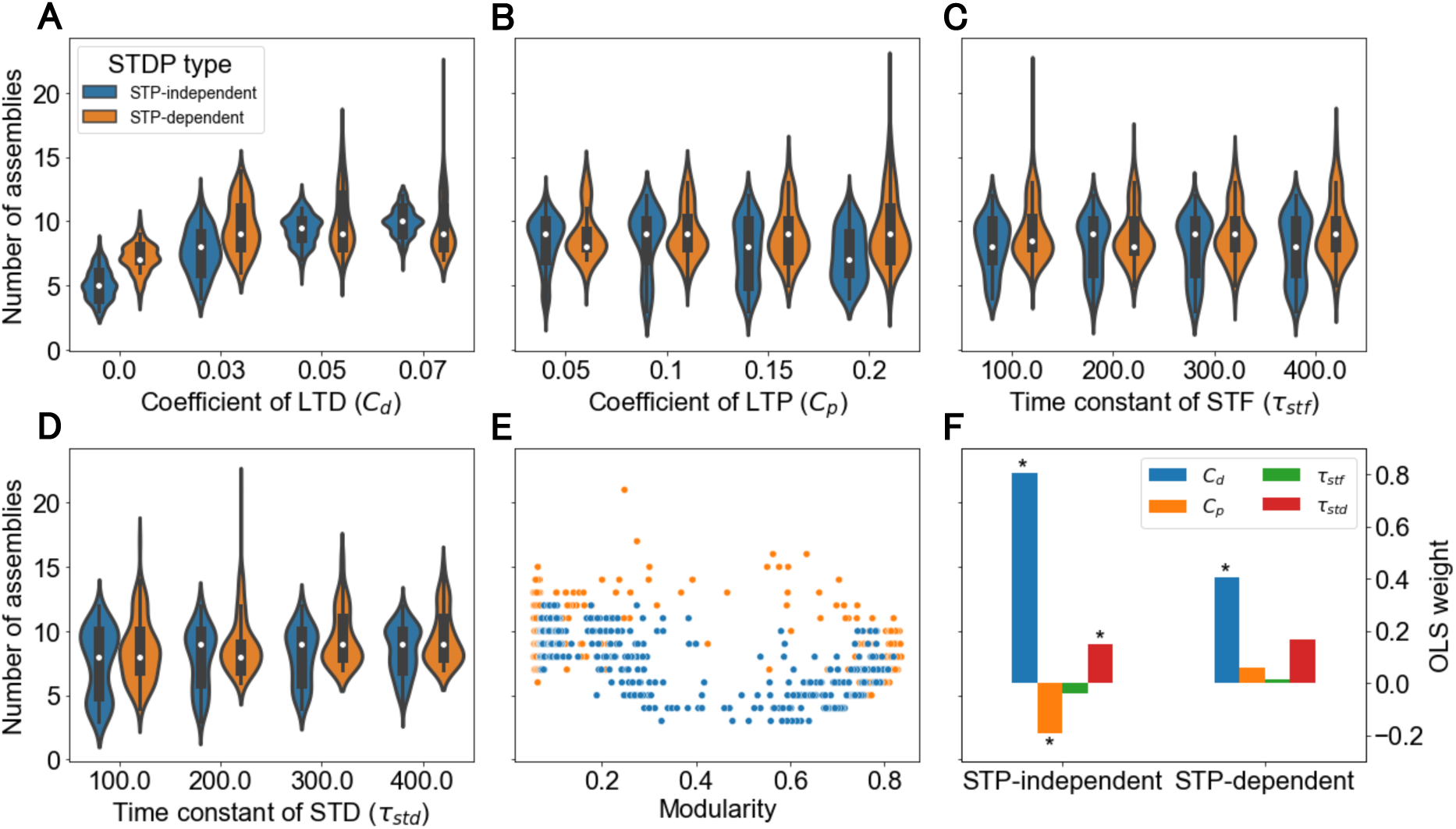
Dependence of the number of spontaneous cell assemblies on plasticity parameters. (A) The coefficient of LTD tends to be positively correlated with the number of spontaneous cell assemblies at equilibrium. By contrast, the coefficient of LTP (B), the time constants of time constants of short-term facilitation (C) and short-term depression (D) are only weakly correlated with the number of cell assemblies at equilibrium. (E) Likewise, there was no obvious linear correlation between the number of cell assemblies and modularity at equilibrium. (F) The coefficients of a multiple linear regression model further confirm that the number of cell assemblies at equilibrium mostly depends on *C_d_*. Stars in (F) indicate the weights of a linear regression model with *p* < 0.01. The effects of the LTP coefficient and STP time constants appear much weaker or non-significant. All the plots are based on 1024 simulations run for 700 s (model time).

### Stability of cell assemblies against representational drift

Recent studies have revealed that, even after an animal has mastered a certain task, the patterns of neural activity associated with the performance of that task, continually change in a process known as “representational drift” (Driscoll et al. 2017; Mau et al. 2020; Rule et al. 2019). Representational drift has been observed in the mouse posterior parietal cortex (PPC), during a task requiring sensorimotor skills, and does not seem to cause the animal to lose performance on that task (Driscoll et al. 2017; Mau et al. 2020; Rule et al. 2019). Similarly, in the hippocampal CA1 region, the spatial mapping of individual neurons shifts over time, even though the environment remains unchanged and familiar to the animal (Driscoll et al. 2017; Mau et al. 2020; Rule et al. 2019). We explored representational drift in our models and show typical results below for our STP-dependent STDP rule.

In our simulations, for many combinations of parameters, the network reaches a point beyond which the number of cell assemblies no longer changes^1^ (Fig. 3A), but, similarly to previous modeling studies (Fauth and van Rossum 2019; Kossio et al. 2021; Manz and Memmesheimer 2023; Tomé et al. 2024; Triplett et al. 2018), the weights of E-E synapses continue to evolve in terms of their self-similarity^2^ (Fig. 3B), cluster labels (Fig. 3C), and the average strength (Fig. 3D). This means that individual connections both between and within cell assemblies continue to evolve, resulting in some neurons occasionally changing their cell assembly “membership”, even after the assembly macrostructure has overall stabilized (Fig. 3C). To further inspect the representational drift of cell assemblies, we visualize how the cell assembly structure evolved in time. Figure 3E shows the instantaneous weight matrices sorted with neurons’ cluster labels obtained at 500 s (i.e. after the cell assemblies reached equilibrium). The weight matrices at later times show progressively stronger off-diagonal elements, implying that some neurons changed their cluster memberships. However, the same weight matrices sorted with cluster labels obtained at the corresponding times do not have visible off-diagonal elements, indicating that the cell assembly macrostructure was mostly preserved over a long period of time (Fig. 3F, S3). Thus, while representational drift does take place under STP-dependent STDP, it is slow enough to preserve the overall assembly structure.

**Figure 3.**
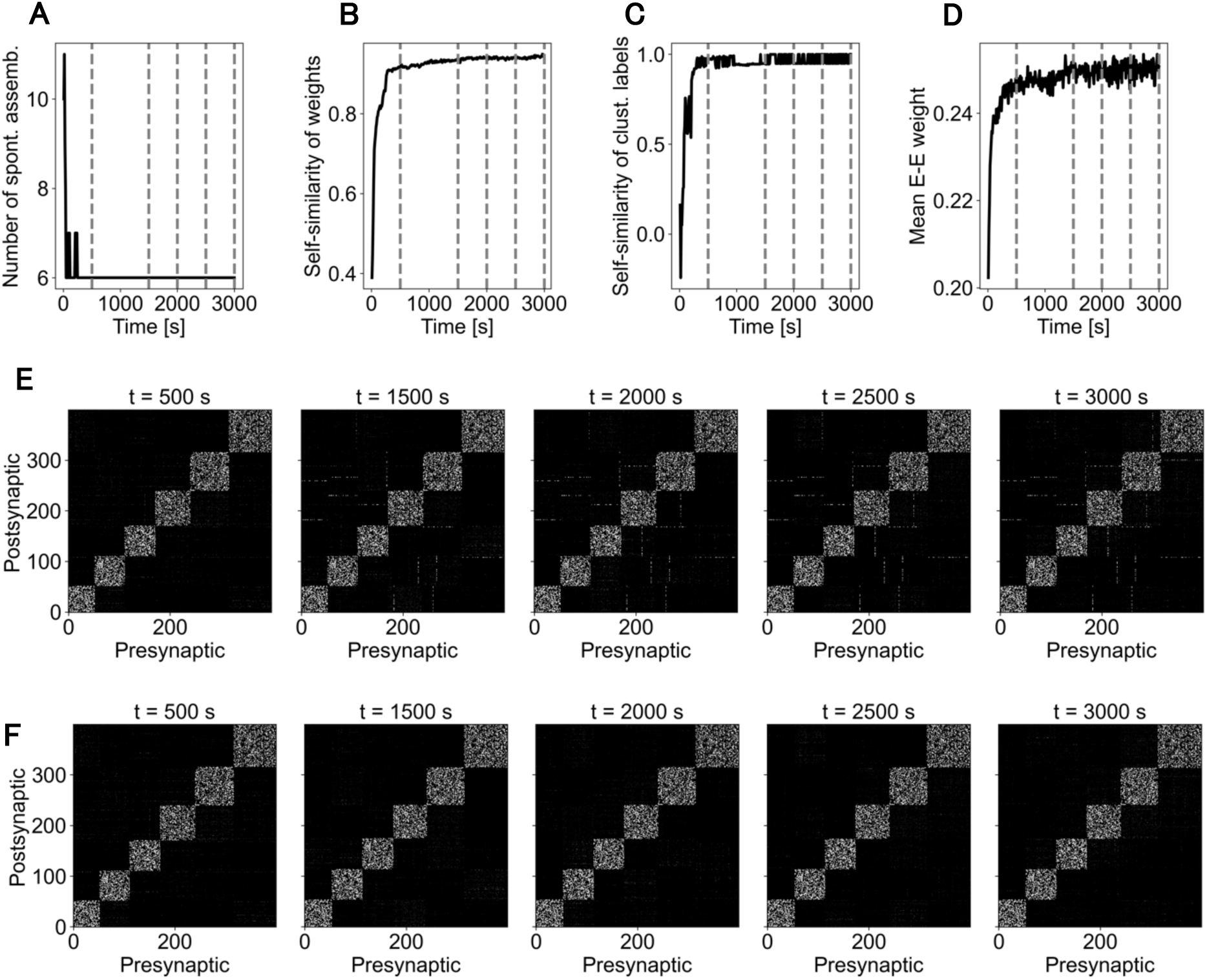
**Representational drift after cell assembly stabilization**. All results are shown for an STP-dependent STDP model. (A) While the number of spontaneous cell assemblies stabilizes at about 500 s, synaptic weights continue to evolve. These changes are reflected by the changing self-similarity of weights (B) and cluster labels (C), as well as continued growth of the mean weight of connections between excitatory neurons (D). Note that the slopes of curves in B, C and D become shallower at around the time at which the number of cell assemblies stabilizes. (E) Weight matrices at times later than 500 s are sorted with the cluster labels calculated at 500 s to visualize an ongoing representational drift (see off-diagonal elements at later times). (F) The same weight matrices sorted with the cluster labels calculated at the corresponding times show the overall stability of cell assembly structure. Vertical grey dashed lines in A-D mark the times at which weight matrices shown in E and F were recorded.

### Regulatory role of astrocytes in spontaneous self-organization of cell assemblies

So far, we have studied the fundamental dynamical features of cell-assembly formation in recurrent network models evolving under symmetric STDP with and without modulation by STP. In the former, STP not only affects excitatory synaptic transmission, but also directly modifies the LTP/LTD amplitude for a given pre-post spike time difference. Now we turn to our central question, the astrocyte-regulation of cell assembly formation (mainly) through the STP-dependence of STDP. Chipman et al. (2021) investigated the role of astrocytic GluN2C NMDA receptors on excitatory synaptic transmission onto hippocampal CA1 pyramidal neurons and found that selective blocking of those receptors reduced the diversity of neurotransmitter release probabilities and, by proxy, weakened long-term potentiation. Mathematical modeling further confirmed a stronger expression of LTP under a diverse, rather than homogeneous, distribution of presynaptic release probabilities.

The fact that astrocytes amplify the expression of LTP raises the question whether and how astrocytes might modulate the emergence and behavior of cell assemblies, including their stability and robustness to perturbations. To address this question, here we indirectly modeled astrocyte effects by manipulating the neurotransmitter release probability at E-E synapses and ran our model in four conditions: with and without astrocyte effects (astro+ and astro-) under STP-dependent or STP-independent STDP. With the latter STDP, astrocytes only modulated excitatory synaptic transmission without affecting long-term synaptic plasticity. Following Chipman et al. (2021), in the astro+ condition, each excitatory neuron’s release probability was obtained by sampling from a gamma distribution with shape and scale parameters set to 2 and 30, respectively, and dividing the sampled values by 130 to ensure they fall within a range 0 to 1 and have a mean of 0.3. In astro-, the release probability was set to 0.3 for all excitatory neurons, exactly matching the mean release probability in the astro+ conditions (Fig. 4A).

**Figure 4.**
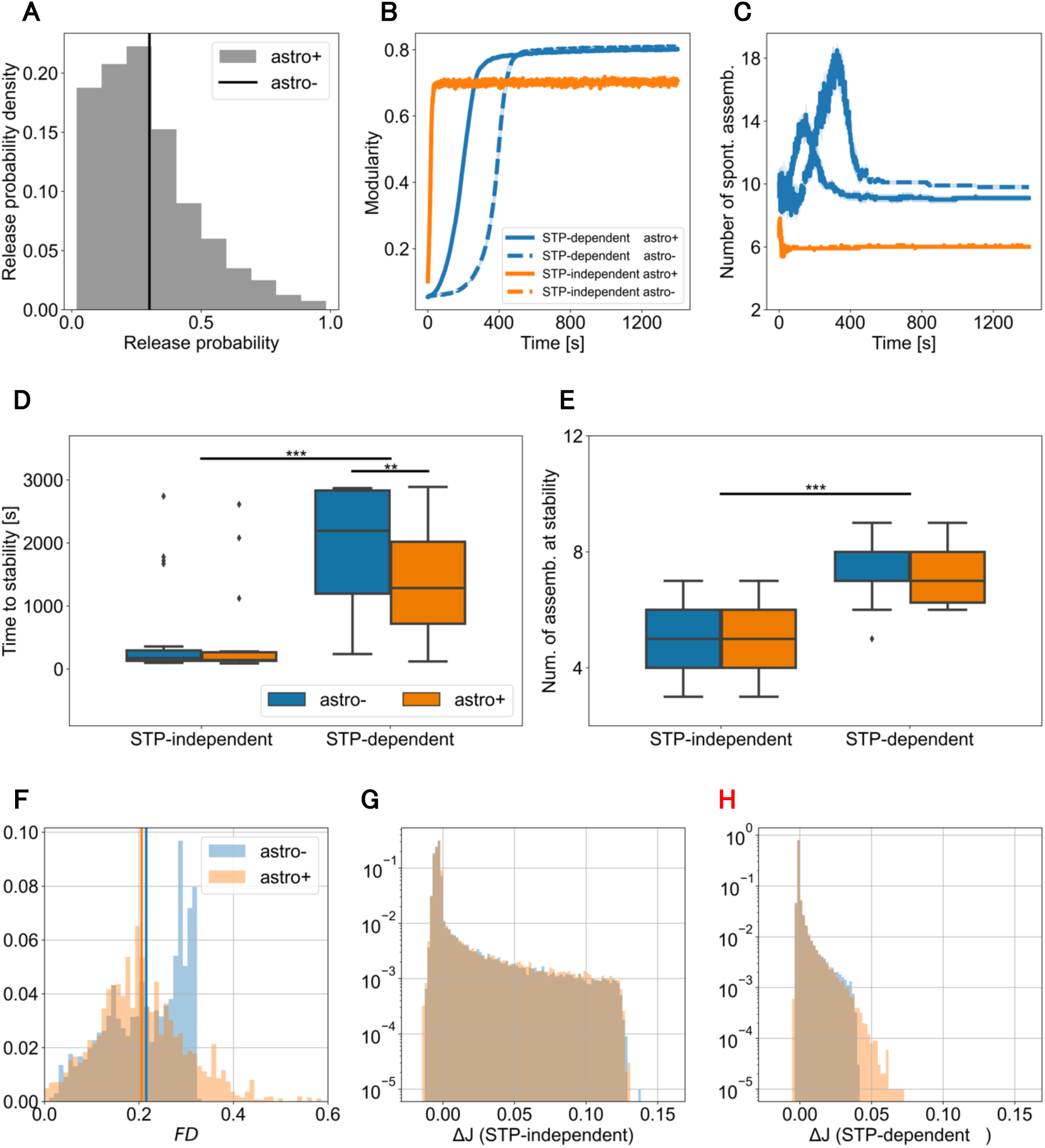
Astrocyte regulation of cell assembly formation under STP-dependent STDP. (A) The distributions of release probabilities in the astro+ and astro-conditions. The time evolution of modularity (B) and number of cell assemblies (C) are shown under the four conditions. Shades in A and B indicate the SEM over 10 simulations with identical parameters but different weight initializations. In B, traces for the astro+ and astro-conditions overlap with each other and cannot be distinguished. Time to stability (D) and the number of stable cell assemblies (E) are compared quantitatively between the four conditions. The distributions of the product *FD* (F), which describes an instantaneous release probability in STP, and the instantaneous weight change under STP-independent (G) and STP-dependent STDP (H) both for the astro+ and astro-conditions.

Our model of astrocyte regulation is certainly an oversimplification of the complex astrocyte-neuron interactions. Nonetheless, it highlights the crucial role astrocyte play in cell assembly formation, and specifically in establishing a trade-off between the robustness and flexibility of cell assemblies. First, self-organization proceeds much faster under STP-independent rather than STP-dependent STDP, and this situation does not change with whether astrocytes regulate STP or not (Fig. 4B). However, the rapid self-organization with STP-independent STDP comes at the cost of lower modularity and number of stable cell assemblies, where the latter becomes about 40% smaller than that of STP-dependent STDP (Fig. 4C). Astrocyte regulation slightly accelerates the self-organization of the weight structure under STP-dependent STDP, reducing the time necessary for cell assembly formation by approximately 50% (Fig. 4B, D). In addition to faster self-organization, astrocyte regulation does not seem to systematically affect the size of the stable cell assemblies (Fig. 4C, E), resulting in roughly the same number of cell assemblies at stability. The accelerated growth of modularity (blue curves in Fig. 4B) is likely due to the longer right tail of the STP factor (Fig. 4F), which directly modulates the weight change due to STP-dependent STDP (compare Fig. 4G and H). Thus, our model suggests that astrocyte regulation accelerates spontaneous self-organization under STP-dependent, but not STP-independent, STDP. These results further illuminate astrocytes’ possible regulatory role for cell assembly dynamics.

### Stimulus-driven network remodeling under STP-dependent STDP

The hippocampus is thought to form novel memories by remodeling its network structure through new experiences. A recent study suggested that increased astrocytic cAMP induces LTP in a neural population to encode a new memory into a long-lasting one (Zhou et al. 2021). In short, astrocyte-induced synaptic plasticity enhances the formation of new memories while it impairs the retention of existing memories. Motivated by these findings, we investigated the flexibility and robustness of the self-organized weight structure, noting that these characteristics pose conflicting demands on the stability of memory traces. To this end, we observed how an external stimulus modifies this structure under the four conditions described above, i.e., with STP-dependent vs. STP-independent STDP, and astro+ vs. astro-conditions. In each of them, we stimulated a random subset of 200 neurons in a fully stabilized network initialized with weights of a model which had been run for 1000 s until stability for 50 s (i.e., from 1050 to 1100 s) and observed how this perturbation changed the structure of cell assemblies.

In all the four conditions, the modularity of cell assemblies (Fig. 5A) and the average E-E weights (Fig. 5B) decreased during the stimulus, but these quantities rapidly returned to the original levels after stimulus termination. Therefore, we could unambiguously define cell assemblies before and after the stimulus. However, the modularity dropped during the stimulus more significantly for STP-independent than STP-dependent STDP. The number of cell assemblies and self-similarity of weight matrices showed contrasting features between the different conditions. Under STP-independent STDP, despite a very strong weight drift during and after the stimulus (Fig. 5C, G), the number of cell assemblies before and after the stimulus remained roughly the same (Fig. 5D).

**Figure 5.**
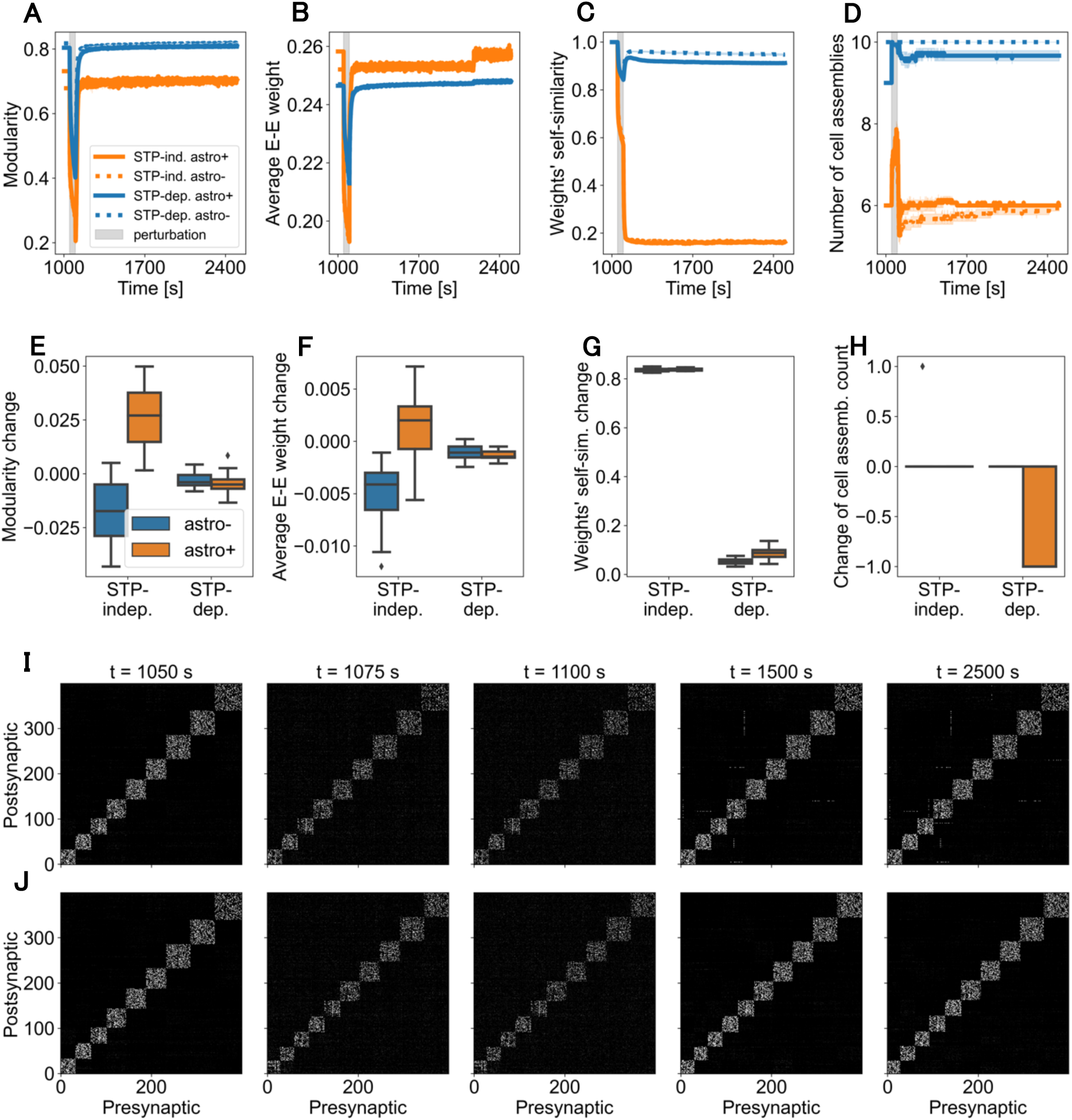
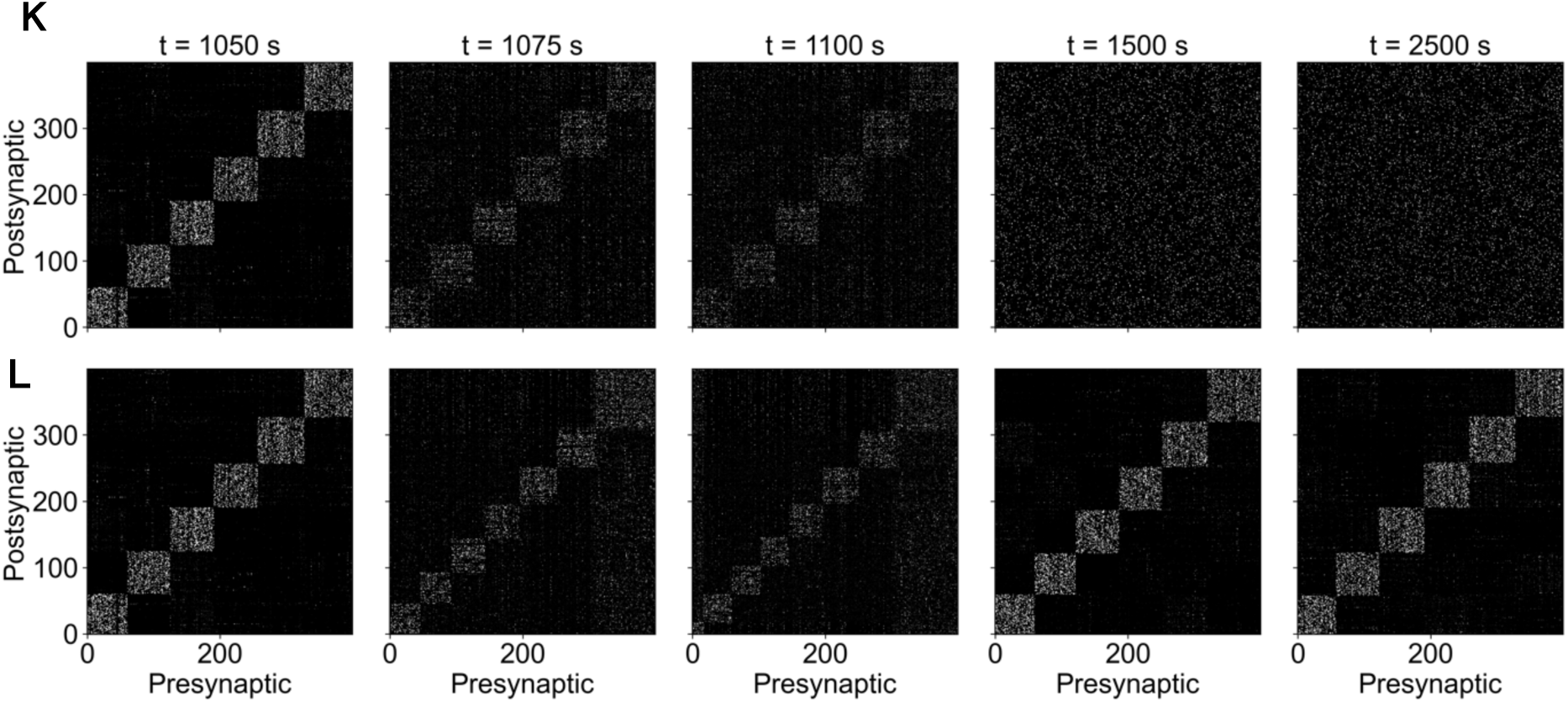
Under STP-dependent STDP astrocytes facilitate stimulus-driven cell assembly reorganization. A 50-second-long stimulus (indicated by a grey vertical bar) applied to randomly chosen 200 neurons applied to a stabilized network causes changes in (A) the modularity, (B) average E-to-E weights, (C) self-similarity of weights, and (D) number of cell assemblies in all the four conditions. Lines and shades in A-D represent the mean and 1 SEM, respectively, over 15 simulations. Self-similarity in C is shown relative to the time step preceding stimulus onset (1050 s). Panels E-H illustrate the differences in the recorded quantities between the beginning and end of the simulation. The E-E weight matrices evolving with STP-dependent STDP are sorted with the cluster indices obtained at (I) the time 1050 and (J) the corresponding times. The former matrices develop off-diagonal elements indicating a slow representational drift. For comparison, weight matrices evolving much faster with STP-independent STDP are sorted with the cluster indices obtained at (K) the time 1050 and (L) the corresponding times. These weight matrices were obtained in the astro+ condition, although the astro-condition yielded essentially the same results.

By contrast, under STP-dependent STDP, the weight drift during the stimulus presentation was much slower (Fig. 5C), and, as a consequence, weight matrices were similar before and after the stimulus. In addition, the number of cell assemblies remained large in both *astro*+ and *astro*-conditions (Fig. 5D). Interestingly, in the *astro+* condition, the stimulus caused a larger change (than in *astro-)* both on the level of weight self-similarity and cell assembly structure: the number of cell assemblies increased from nine to ten during the stimulus presentation and stayed at an average of about 9.5 (Fig. 5D). Figure 5E-H show the differences in those quantities between the beginning and end of the simulation. The greater sensitivity of cell assembly structure in the *astro+* than astro-condition under STP-dependent STDP (Fig. 5C) is consistent with the accelerated self-organization we reported in the previous section, suggesting that astrocyte regulation of synaptic plasticity might facilitate learning on exposure to new stimuli.

We visually inspect the time evolution of weight matrices during and after the stimulus offset in both STP-dependent (Fig. 5I and J) and STP-independent STDP (Fig. 5K and L) in the astro+ condition. In Fig. 5I and K, the weight matrices are sorted with neurons’ cluster labels obtained at the stimulus onset (i.e., the time 1050 s), whereas in Fig. 5J and 5L they are sorted with cluster labels obtained at the corresponding times. These figures demonstrate clearly that the stimulus modified the cell assembly structure for both STP-dependent and independent learning rules. However, under STP-dependent STDP, the stimulus left the cell assembly structure largely unchanged (Fig. 5I and 5J), under STP-independent STDP we observed a significant disruption of the pre-existing cell assemblies (Fig. 5K and 5L), which took a relatively long time to recover. In addition, representational drift, indicated by off-diagonal elements emerging after the stimulus termination, occurred much slower with STP-dependent than STP-independent STDP (compare Fig. 5J and Fig. 5K).

In sum, STP-dependent STDP, but not STP-independent one, can balance the robustness and flexible remodeling of cell assembly structure. Preserving the learned cell assemblies against novel sensory inputs is thought to be crucial for episodic memory processing by the hippocampus. Essentially the same results were obtained in a wider range of stimulus parameters (Fig. S1).

### Stability of learned cell assemblies under STP-dependent STDP

A recurrent neural network with a finite size can generally store only a finite number of stable cell assemblies (Brunel 2016; Gardner 1988; Khona and Fiete 2022), where a typical example is associative memory models (Amit, Gutfreund, and Sompolinsky 1985; Burns and Fukai 2023; Chapeton et al. 2012; Hopfield 1982; Krotov 2016, 2023). The limitation of storage capacity of the present model (with 500 excitatory neurons) follows from the fact that it always spontaneously self-organizes a similar number of cell assemblies. Below, we analyzed the storage capacity of the network in encoding cell assemblies formed in response to structured stimulation involving a certain number of patterns.

We stimulated the network for 150 s with *M* ∈ {2, 4, 6, 8, 10, 12, 14, 16, 18, 20} non-overlapping patterns of *S* ∈ {200, 100, 80, 66, 50, 40, 33, 28, 25, 22, 20} neurons respectively (see Materials and methods). We presented these patterns in random order and observed the stability of the resulting cell assemblies over 240 s after stimulus termination. As in the previous sections, we examined the stability in the four different conditions. Figure 6 shows the results for *M* = 8 to exemplify a specific case where the stimulus-induced assembly structure remains permanently stable (for STP-dependent STDP). Results for other values of *M* are provided in Fig. S2.

**Figure 6.**
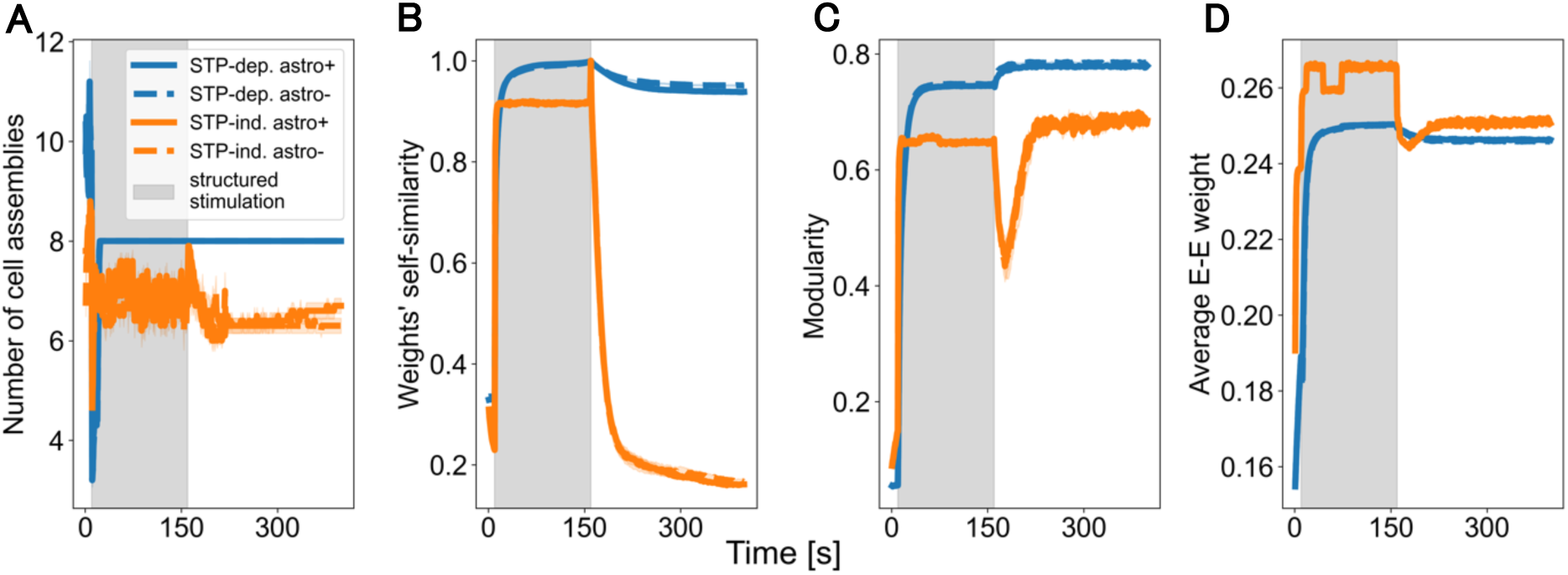
Cell assemblies resulting from structured stimulation. Here, *M* = 8. (A) STP-dependent STDP (blue) could encode a larger number of more stable cell assemblies than STP-independent one (orange). Increased stability is likely due to a very slow weight drift (B), relatively high modularity (C) and stable average weight (D).

In general, the learned cell assembly structure is more stable under STP-dependent STDP (Fig. 6, blue lines) than under STP-independent STDP (Fig. 6, orange lines). This is convincingly indicated by higher modularity (Fig. 6C) and is likely due to a very slow weight drift under the former plasticity rule (Fig. 6B). In general, cell assemblies can be considered distinct and stable if the modularity is greater than about 0.3. The stability of learned cell assemblies was not significantly affected by the presence of astrocyte regulation. We note that cell assemblies appeared to be stable only for *M* ∈ {6, 8, 10}. In other cases, irrespective of the number of patterns enforced by an external stimulus, the network eventually evolves into a stable state with this range of cell assembly numbers (Fig. S2). This suggests a relationship between the network size and number of stable cell assemblies that can exist in the network under a specific plasticity rule.

### Under STP-dependent STDP, astrocytes marginally facilitate the orthogonalization of learned cell assemblies

As we have seen in the preceding section, the network’s weight structure evolves towards maximum modularity at which the spontaneous activity becomes orthogonal: while one cell assembly is active, the rest of the excitatory population is suppressed via the inhibitory network. How do STP dependence of STDP and astrocyte regulation influence this property?

To answer this question, we stimulated 8 subsets of neurons overlapping in space by 50% for 200 s (Fig. 7A). We conducted this simulation for STP-dependent STDP in the astro+ condition. At the end of stimulation, this resulted in cell assemblies having significant mutual overlaps (Fig. 7B). However, such a cell assembly structure was fragile and eventually settled into a more stable “preferred” structure with maximum modularity. Figure 7C displays the distributions of cofiring probabilities of neuron pairs belonging to the same and different cell assemblies (see Materials and methods for details). Co-firing probabilities are high in within-assembly pairs and low in between-assembly pairs, and the two distributions do not overlap significantly in the “preferred” network structure (Fig. 7C, right). This implies that neurons belonging to different assemblies rarely co-fire, demonstrating that the cell assemblies are almost completely orthogonalized in the preferred structure.

**Figure 7.**
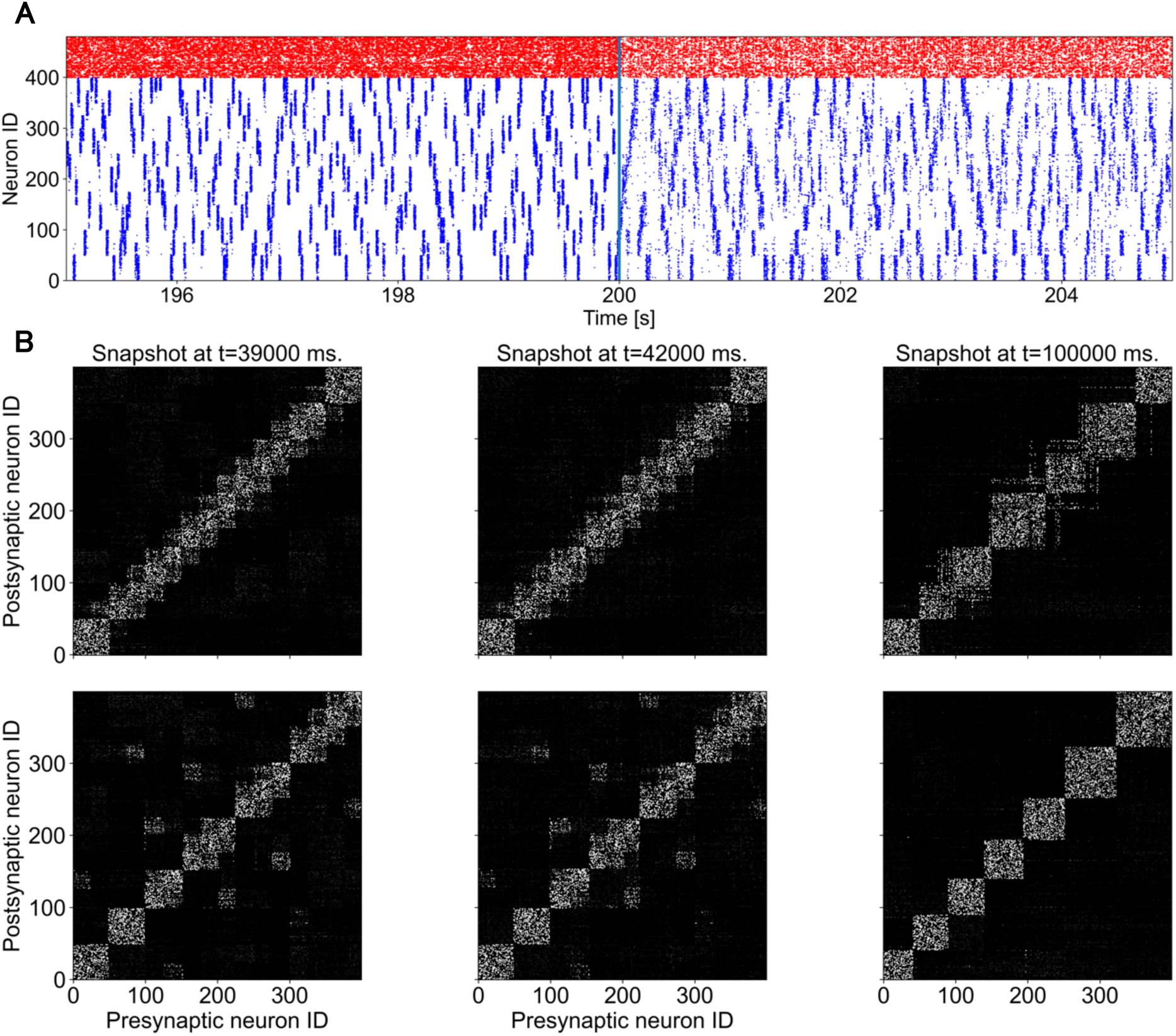
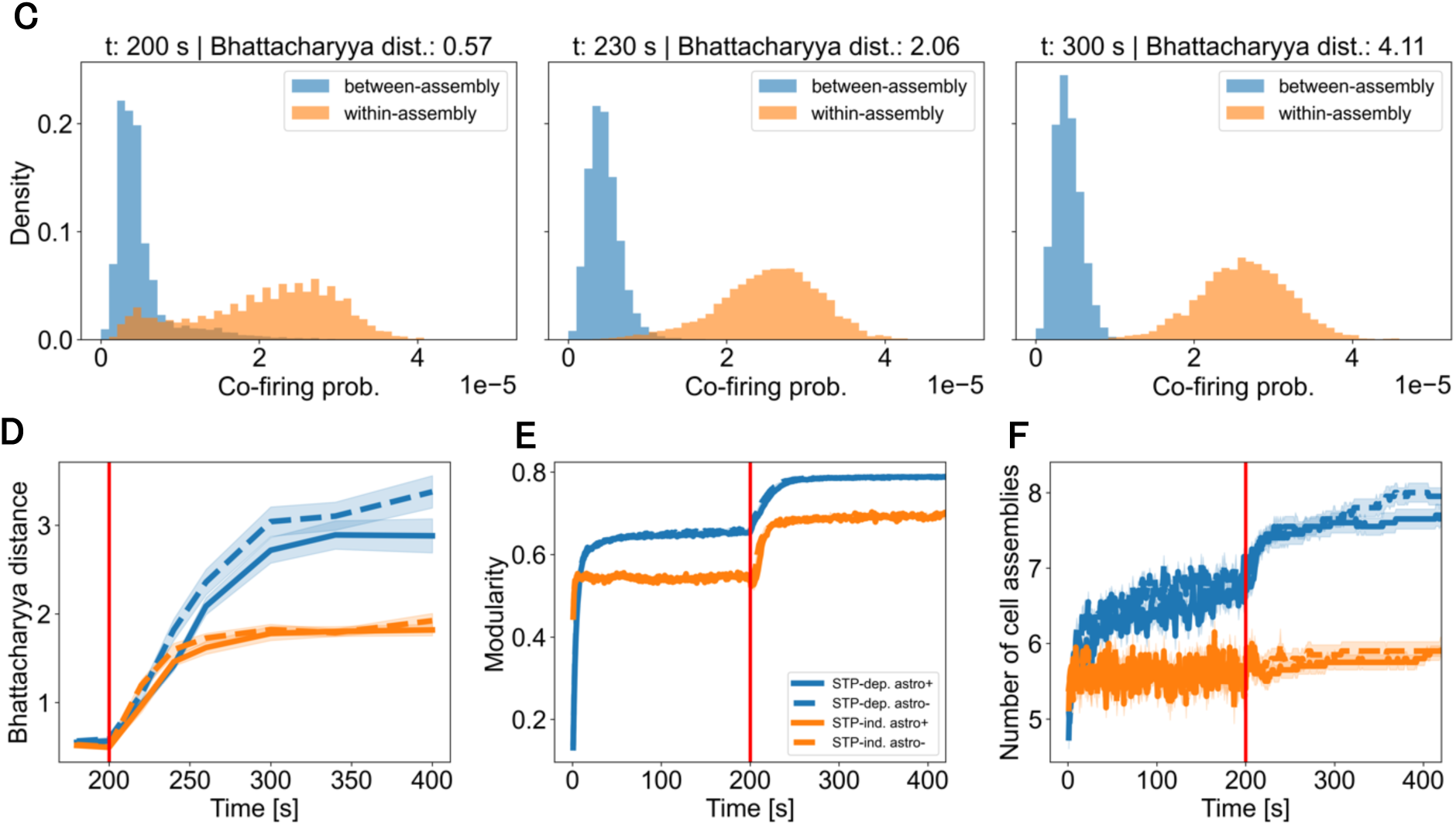
Orthogonalization of learned cell assemblies. In panels A-C, results are shown for STP-dependent STDP in the astro+ condition. In panels D-F, we compare results across all the four conditions. (A) Spike rasters of excitatory (blue) and inhibitory (red) neurons, illustrating activity before and after the stimulus offset (vertical line). Eight overlapping subsets of neurons were stimulated for 200 s. (B) The evolution of weight matrices post-stimulus is displayed, sorted according to the cluster indices obtained at 200 s (top) and the corresponding subsequent times (bottom). Notably, the bottom rightmost weight matrix lacks off-diagonal elements, highlighting the achieved orthogonalization. (C) Displays the distributions of co-firing probabilities for neuron pairs within the same (orange) and across different (blue) cell assemblies at three post-stimulus time points. The time evolution is shown for the distances between the co-firing probabilities (D), modularity (E), and the instantaneous number of cell assemblies (F).

We further studied the orthogonalization process in all four conditions (STP-independent and STP-dependent, *astro*+ and *astro*-), paying attention to how astrocyte regulation influences its time course and final network state. The Bhattachryya distance (Bhattacharyya 1943) between the co-firing probability distributions of within- and between-assembly pairs gradually increased in all the four conditions, and the maximum separation was much larger for STP-dependent than STP-independent STDP (Fig. 7D). Astrocyte regulation slightly suppressed the maximum separation for the former learning rule. The network gained higher maximum modularity (Fig. 7E) and number of cell assemblies (Fig. 7F) under STP-dependent than STP-independent STDP, while cell assemblies stabilized slightly earlier under the latter learning rule. These results are overall consistent with the simulation results shown previously. In sum, the orthogonalization occurs much more strongly under STP-dependent than STP-independent STDP, but this process does not significantly rely on astrocyte regulation.

### Cell assembly stability analysis

Why is cell assembly structure stable mostly with 6 – 10 assemblies for our chosen network and plasticity parameters? A mean-field analysis of the storage capacity of recurrent networks, previously for a similar STP-dependent symmetric STDP without astrocyte regulation (Hiratani and Fukai 2014), showed that multiple cell assemblies can be stably maintained only for relatively small release probabilities within a certain value range. In such a case, the activation of each cell assembly has a sufficiently long lifetime, enabling stable spontaneous re-activation of the cell assemblies. Strong cell assemblies remain active as long as their constituent neurons, on average, have a sufficient amount of neurotransmitter to maintain active firing of downstream neurons and, through feedforward inhibition, suppress activity of the rest of the network. The time it takes a cell assembly to deplete its supply of neurotransmitters – given the same firing rate and average connection weight network-wide – is proportional to its size: larger cell assemblies will tend to fire longer than smaller ones (Fig. 8C). However, if a cell assembly is too large, such that it fires longer than the positive part of the STDP function, it will be unstable and likely to break up into smaller cell assemblies. This is because neurons that fired at its beginning and end will be less strongly wired to each other than to the neurons that fired in the middle. On the other hand, small cell assemblies, such that more than one of them fit within the positive part of the STDP function, will tend to merge into larger ones by the same principle. In addition, given an upper bound on the firing rate and connection strength, small cell assemblies cannot produce enough excitation in the inhibitory population to suppress activity of the other cell assemblies and therefore will not be able to maintain their identity as a cell assembly. For our chosen network configuration and STDP and STP parameters, the stable number of cell assemblies is between 6 and 10. Short-term plasticity appears to be the main determinant of cell assembly lifetime: to deplete the neurotransmitter supply, each neuron in a cell assembly needs to fire an average of 4-5 spikes within a short period of time (Fig. 8A).

**Figure 8.**
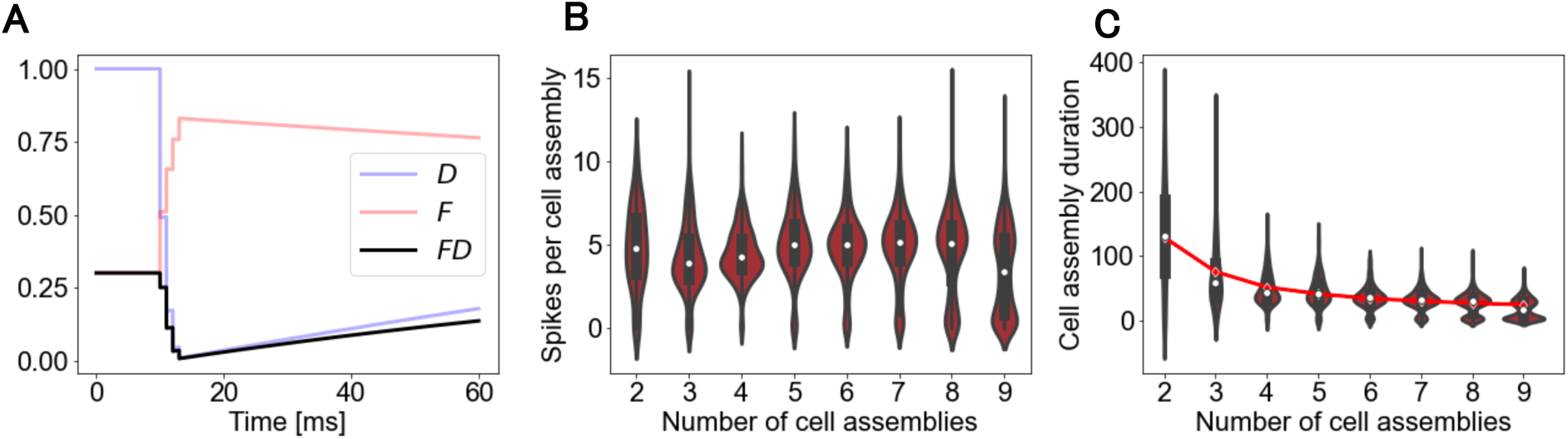
Dependence of the network’s capacity on STP. (A) Responses of STP variables a spike burst. About four spikes are needed on average to deplete the neurotransmitter available at the presynaptic terminal. (B) Observed number of spikes per cell assembly as a function of the number of cell assemblies in the network. The numbers approximately coincide with the number of spikes necessary for depleting presynaptic neurotransmitters shown in A. (C) The duration of cell assembly activations (refer to Material and Methods) as a function of the number of cell assemblies in the network. The predicted average duration *L* of a cell assembly’s activity is also shown (red line: see the main text).

This is roughly consistent with the observed number of spikes in cell assemblies of different sizes (Fig. 8B). In fact, it is possible to predict the duration of cell assemblies using the following simple model: *L* = *n*/(ν*M*), where *L* is the expected duration of a cell assembly (in seconds), *n* is the average number of spikes it takes a neuron to deplete its neurotransmitter, ν is the average firing rate of the excitatory population (which in our simulations was around 20 Hz), and *M* is the number of cell assemblies (Fig. 8C).

## Discussion

We investigated the activity-dependent formation, maintenance, and modulation of cell assemblies with and without structured input in a recurrent network model that mimics the hippocampal area CA3. We compared these processes between four variants of symmetric STDP, which take the STP-dependence of STDP and astrocyte regulation of STP into account. Here, we implicitly modeled the effect of astrocytes on the evolution of network structure by introducing large variability into the neurotransmitter release probability. Our model showed several advantages of STP-dependent STDP over more conventional STP-independent one, particularly, for balancing conflicting computational demands, i.e., the robustness and flexibility of the cell assembly structure.

Consistent with our previous study with the conventional symmetric STDP (Hiratani and Fukai 2014), our simulations showed that the spontaneous emergence of cell assemblies can occur in a recurrent network when symmetric STDP is inherently coupled with STP. This modified STDP hypothesizes that the amplitude of synaptic weight modifications is proportional to the amount of releasable neurotransmitter at presynaptic terminals. The learning rule was previously proposed to facilitate sequence learning with time-symmetric STDP (Haga and Fukai 2018). In this study, we further explored whether and how cell assemblies can spontaneously emerge, drift, and are modifiable by external stimuli under STP-dependent symmetric STDP. STP-dependent learning rules self-organize more cell assemblies with a more distinct clustering structure than STP-independent rules (Fig. 4). In other words, the STP-dependence of the learning rule increases the stability of learned cell assemblies and the capacity of the recurrent neural network for storing memory traces. However, this costs a slower speed of self-organization and, for the same reason, remodeling existing cell assemblies also takes longer under STP-dependent than STP-independent STDP. Nonetheless, these dynamical features also bring a positive side in that these structures become more stable against representational drift, maintaining their self-similarity longer across time (Fig. 5, 6, S3).

Intriguingly, the astrocyte regulation of STP mitigates the slow-down of the self-organization process by about 50% without lowering the total number of cell assemblies (Fig. 4). In our model, these are the most computationally beneficial effects of astrocyte regulation on cell assembly formation. This effect of the astrocyte regulation accelerates the speed slightly at which the cell assembly structure returns to a stable configuration allowed by the storage capacity after the cessation of external stimuli (Fig. 5). We could not check whether the cell assembly number returns to the one before the stimuli due to a practical limitation on the simulation time. However, we speculate this should be the case as the weights’ similarity monotonically (but very slowly) decays after stimulus cessation.

In contrast, our results revealed that the astrocyte regulation of STP has no noticeable effects on the cell assembly formation and remodeling if STDP is uncoupled with STP. Even in such a case, synaptic transmissions are regulated by STP in the recurrent network, and therefore the blockade of astrocyte NMDARs influences neuronal dynamics. Our model suggests that astrocyte regulation affects synaptic learning only if STP directly modulates the efficacy of synaptic plasticity. Thus, the computational effects of astrocyte regulation may significantly vary with specific features of synaptic plasticity rules.

Our model supports the hypothesis that astrocytes contribute to neuronal information processing in the brain much more significantly than previously considered. Astrocytes have long been thought to be crucial solely or mainly in the energy supply to neurons. However, accumulating evidence shows the active participation of these cells in a broad range of neural information processing (Chen et al. 2023; Lyon and Allen 2022). For instance, the roles of astrocytes include the enhancement of neuronal synchrony (Fellin et al. 2004), the modulation of synaptic transmissions (Araque et al. 2014; Haydon and Carmignoto 2006), the activity-dependent elimination of synapses (Lee et al. 2021), the regulation of reward signaling pathway (Corkrum et al. 2020), motor learning (Delepine et al. 2023), and the formation, maintenance, and modulation of various types of long-term memory (Akter et al. 2023; Hösli et al. 2022; Navarrete et al. 2019; Sharma et al. 2023; Sun et al. 2024; Vignoli et al. 2021; Zhang et al. 2021; Zhou et al. 2021). Furthermore, impairments in astrocyte regulation can cause a deficit in brain functions (Richetin et al. 2020; Zimmer et al. 2024). Our results suggest additional computational benefits of STP-dependent STDP and highlight a previously unknown regulatory function of astrocytes in memory formation.

## Materials and Methods

We used a recurrent network of binary neurons adapted from (Hiratani and Fukai 2014) to model a small (based on volumetric cell density counts reported in Attili et al. (2019), approximately 0.025 mm^3^) portion of the CA3 area of the hippocampus. Briefly, the network consisted of 2500 excitatory and 500 inhibitory binary neurons, connected with a fixed probability (see Table 1 for the full list of parameters). Each neuron was updated according to:

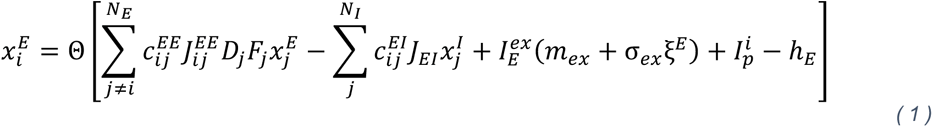

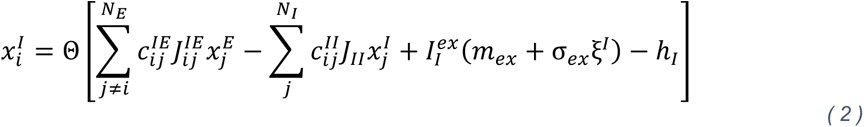

where Θ is the Heaviside step function, N*_E_* and *N_I_* are the number of excitatory and inhibitory neurons in the network, respectively. 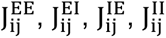 and 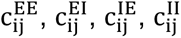 are weights and binary connections for EE, EI, EI, and II synapses, respectively. If there is no connection from neuron *j* to neuron *i*, *c_ij_*_’_ is 0 and 1 otherwise. 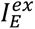 and 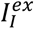 are externally injected random Gaussian noise parameterized by mean *m_ex_* and variance σ*_ex_* and scaled by constants 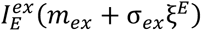 and 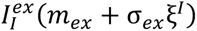 For excitatory and inhibitory populations, respectively. The state of a neuron is set to 1 or 0 depending on whether its membrane potential exceeds the threshold (ℎ*_E_* or ℎ*_I_*) or not.

The original model from (Hiratani and Fukai 2014) was extended in two ways. First, instead of STD-only model of short-term plasticity we will use the Tsodyks-Markram model (Markram, Wang, and Tsodyks 1998), which besides STD also incorporates short-term facilitation (STF):

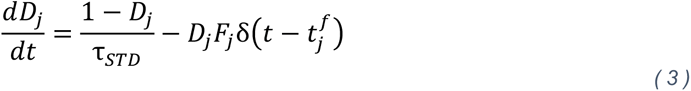

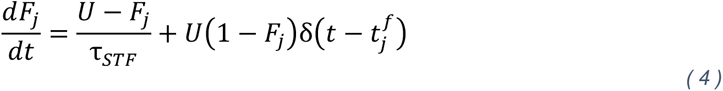

where *D_j_* and *F_j_* are STD and STF terms associated with the *j*-th neuron, *τ_STD_* and *τ_STF_* are their respective time constants, δ is the delta function and *t*_j_*^f^* is a spike time of the *j*-th neuron, and *U* is a fixed free parameter in the range from 0 to 1, modeling the baseline release probability.

This extension is justified by the possibility that, on a short time scale, STF may increase neuronal excitability which is believed to underlie associative memory. Second, the LTP/LTD was modeled not only as a function of temporal difference between a presynaptic and postsynaptic spike, but also the availability of neurotransmitter (STP). To that end, we incorporated a multiplicative STP term into the STDP rule as in (Haga and Fukai 2018):

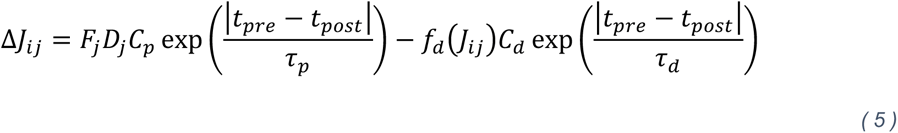

where Δ*J_ij_* is the weight change of the connection between the *j*-th presynaptic and *i*-the postsynaptic neuron, *t_pre_*_-_ and *t_post_* are the spike times of the presynaptic and postsynaptic neurons, respectively, and *C_p_*, *C_d_*, τ*_p_* and τ*_d_* are the STDP coefficients and time constants of potentiation and depression, respectively, and *f_d_*(**J*_ij_*) = ln(1 + α*J_ij_*/*J_EE_*)/ ln(1 + α). A spike is considered to have occurred when the neuron’s state changes from 0 to 1. Eq. 5 is identical to Eq. 3 in (Hiratani and Fukai 2014) except that synaptic weight change due to STDP is now proportional to the amount neurotransmitter available. The rationale for this modification is two-fold. First, experiments suggest that, at least in visual cortex, STDP is attenuated by short-term depression (Froemke et al. 2006). Second, it was demonstrated (Haga and Fukai 2018) that goal-directed sequence learning occurs in a CA3 model only under a symmetric STDP, observed in the rat hippocampus (Mishra et al. 2016). modulated by STP. To prevent the weights from growing without bound, the model included homeostatic plasticity and weight normalization as described in (Hiratani and Fukai 2014).

### Modeling the influence of astrocytic NMDA

In the astro+ condition (see main text), we model the presence of astrocytes with their non-blocked NMDA receptors by sampling *U* (once for each neuron at the beginning of the simulation) from a gamma distribution with shape and scale parameters set to 2 and 30, respectively. Before plugging into Eq. 4, each sampled value of *U* is scaled by diving by 130, values that do not satisfy the constraint 0 < *U* < 1 are rejected. In astro-, which models the condition in which astrocytic NMDA receptors are blocked, the *U* in Eq. 4 is set to the mean of the distribution of *U* = 0.3.

### Perturbation and stimulation

Patterned stimulation (and also perturbation of existing cell assemblies) was achieved as follows. Assume that we stimulate a set *S* ⊂ *N_E_* of excitatory neurons, such that |S| < |*N*_/_|, for *T* timesteps. The subset *S* is chosen once at the beginning of the stimulation. By design, only a very small subset *X* ⊂ *N*, such that |*X*| << |*N*|, of neurons in the network is chosen for update. At each time step *t* ∈ {1 + *d*, …, *T* + *d*}, where *d* is the time at which stimulation starts, ∀*i* ∈ *X* ⋂ *S* we set the membrane potential of the *i*-th neuron to a suprathreshold value.

All the perturbations in Fig. 5 were applied to an already stabilized network, that is a network whose weights and connections at *t* = 0 were initialized randomly according to parameters listed in Table 2 and that was allowed to run without structured stimulation for 1000 seconds of model time.

### Computing the distribution of cofiring probabilities

To compute cofiring probabilities around a particular time in the spike recording 𝒟 = {(*i*, *t*) | *i* ∈ {1, 2, …, *N*}, *t* ∈ {0, Δ*t*, 2Δ*t*, …, *T*}, where Δ*t* is the time step of the simulation, we consider windows of spiking activity 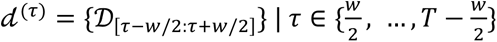 . For the purpose of computing cofiring probabilities, we set *w* = 6000 ms. Then we partition *d*^(*τ*)^ into a set of *K* smaller overlapping windows 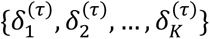, such that 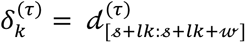, where 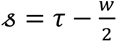 is the left boundary, *l* = 5 ms is the stride and 𝓌 = 20 ms is the width of the each 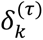. Then we create a zero-filled matrix 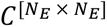 iterating over all *k* ∈ 1, …, *K*, we increment 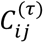 if within 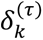 both the *i* -th and *j* -th excitatory neurons spiked at least once. Finally, we normalize the spike counts to get the joint distribution of cofiring probabilities *P*^(*τ*)^ = *C*^(*τ*)^/∑*C*^(*τ*)^. Having cell assembly labels for each neuron at each *τ* allows us to build two probability distributions, one for cofiring probabilities within cell assemblies and the other for cofiring probabilities between cell assemblies. Formally, 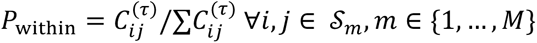, where *S_m_* is a set of neuron indices belonging to the *m*-th cell assembly. Conversely, 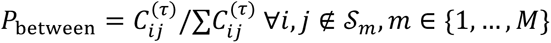.

### Computing the duration of cell assemblies

To determine the duration of cell assemblies, we partition the spike data 𝒟 = {(*i*, *t*, *m*) | *i* ∈ {1, 2, …, *N*}, *t* ∈ {0, δ*t*, 2δ*t*, …, *T*}, *m* ∈ {1, …, *M*} into non-overlapping windows {𝒟_[*t*:*t*+*w*]_}, where *w* = 4 ms, and δ = 0.01 ms. The cell assembly labels, *m* ∈ {1, …, *M*}, are determined with the Louvain clustering algorithm (Blondel et al. 2008) available in the scikit-network package for Python (Bonald et al. 2020). For each window {𝒟_[*t*:*t*+*w*]_}, we count the number of spikes in each cell assembly. The cell assembly, which emitted the most spikes is considered active withing the given window: 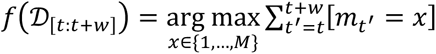. Having determined the most active cell assembly within each window, we can also determine the length of the periods, during which each of the M cell assemblies were active, and build a distribution of these lengths for each 𝒟.

## Supplementary Figures

**Figure S1.**
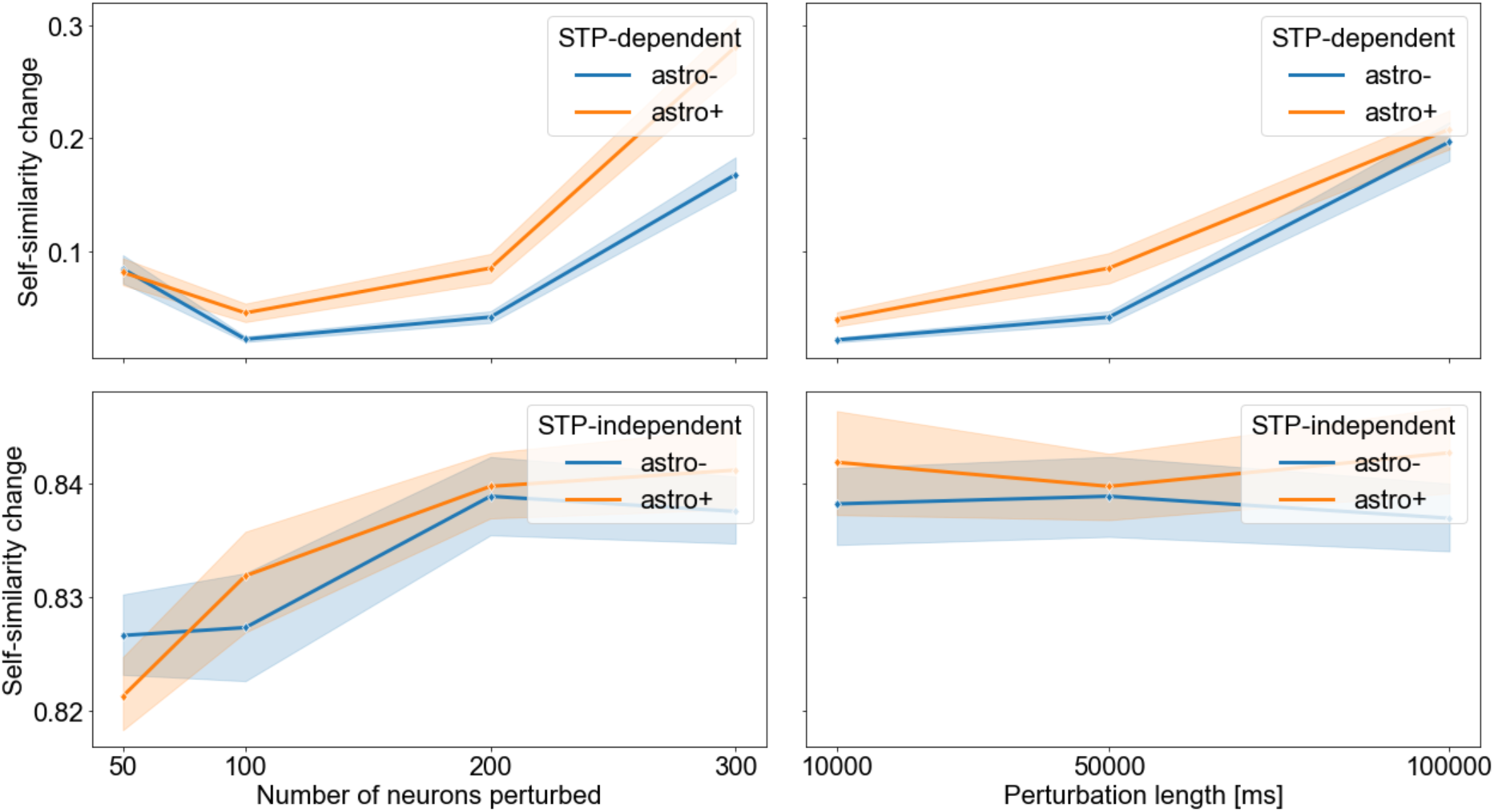
Under STP-dependent STDP self-organized cell assemblies respond more readily to perturbations. Here, in model with stabilized self-organized cell assembly structure, we perturbed 50, 10, 200 and 300 neurons for 10, 50 and 100 s. The perturbations resulted in greater weight self-similarity changes under STP-dependent STDP (astro+).

**Figure S2.**
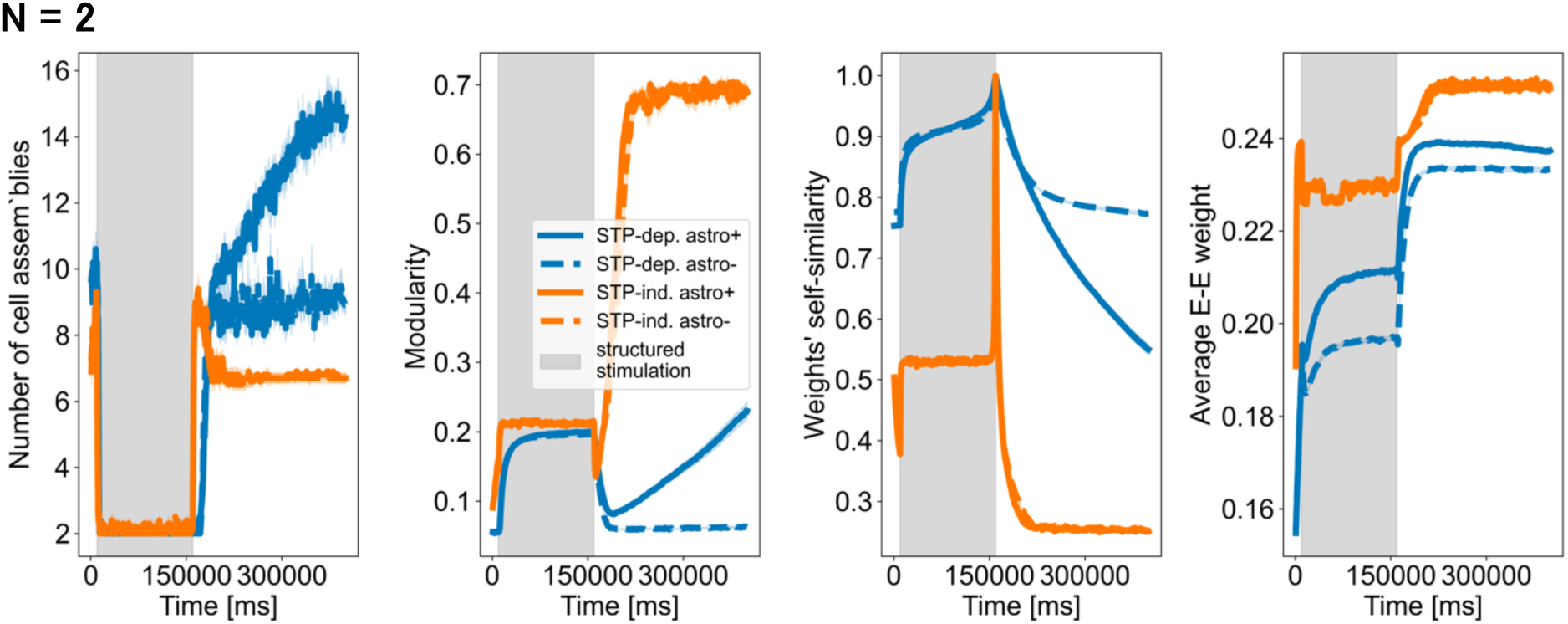
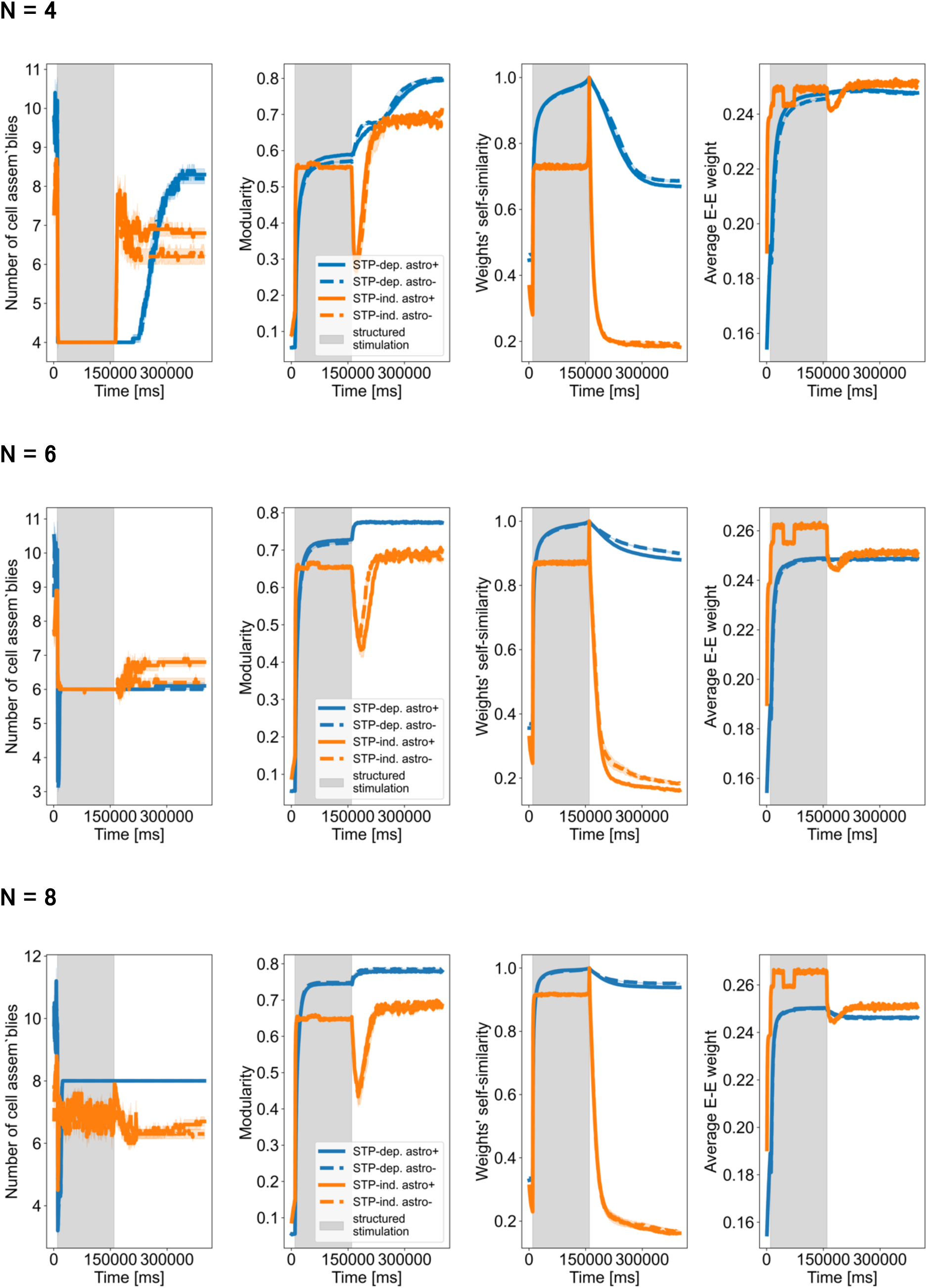
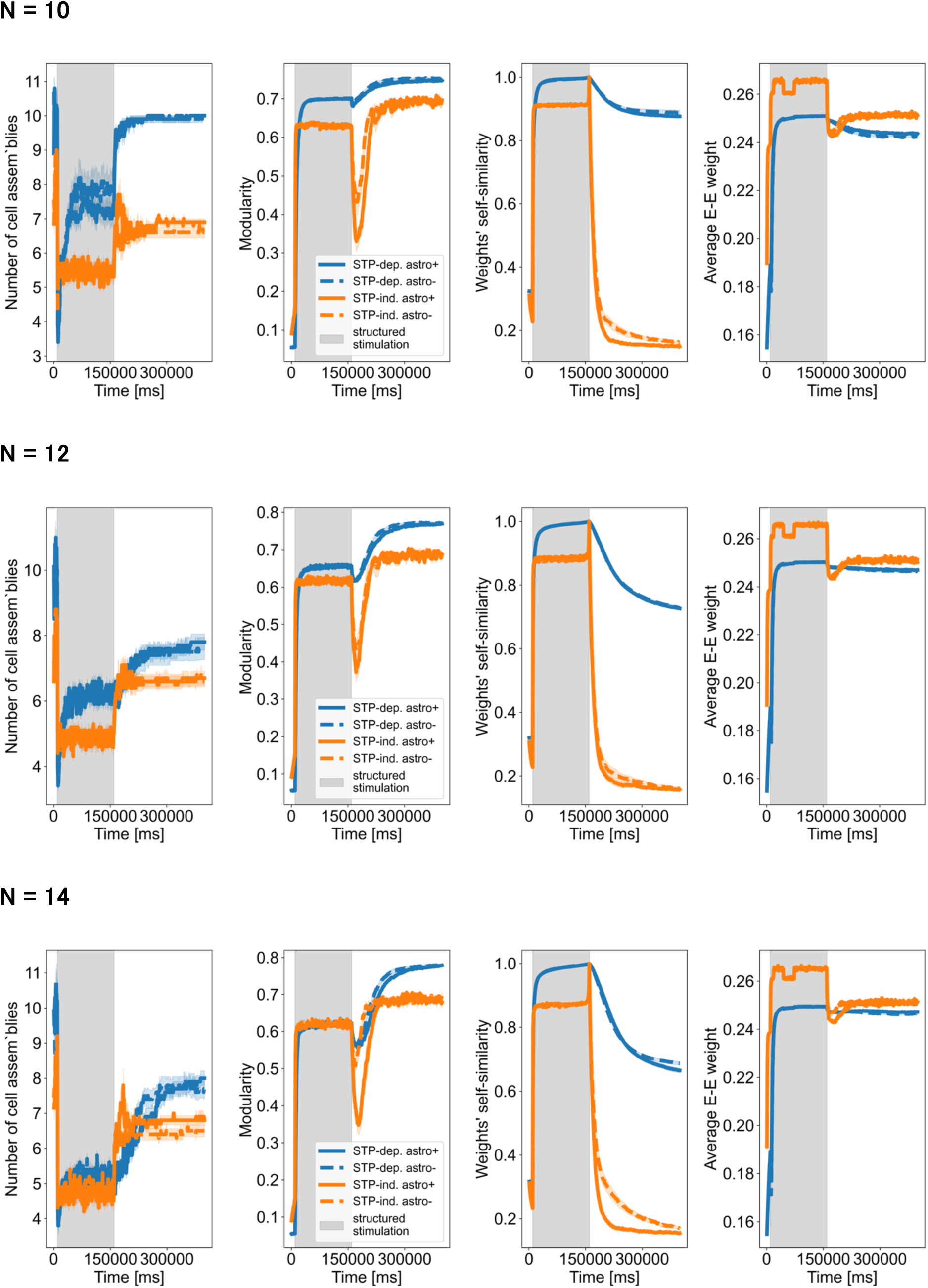
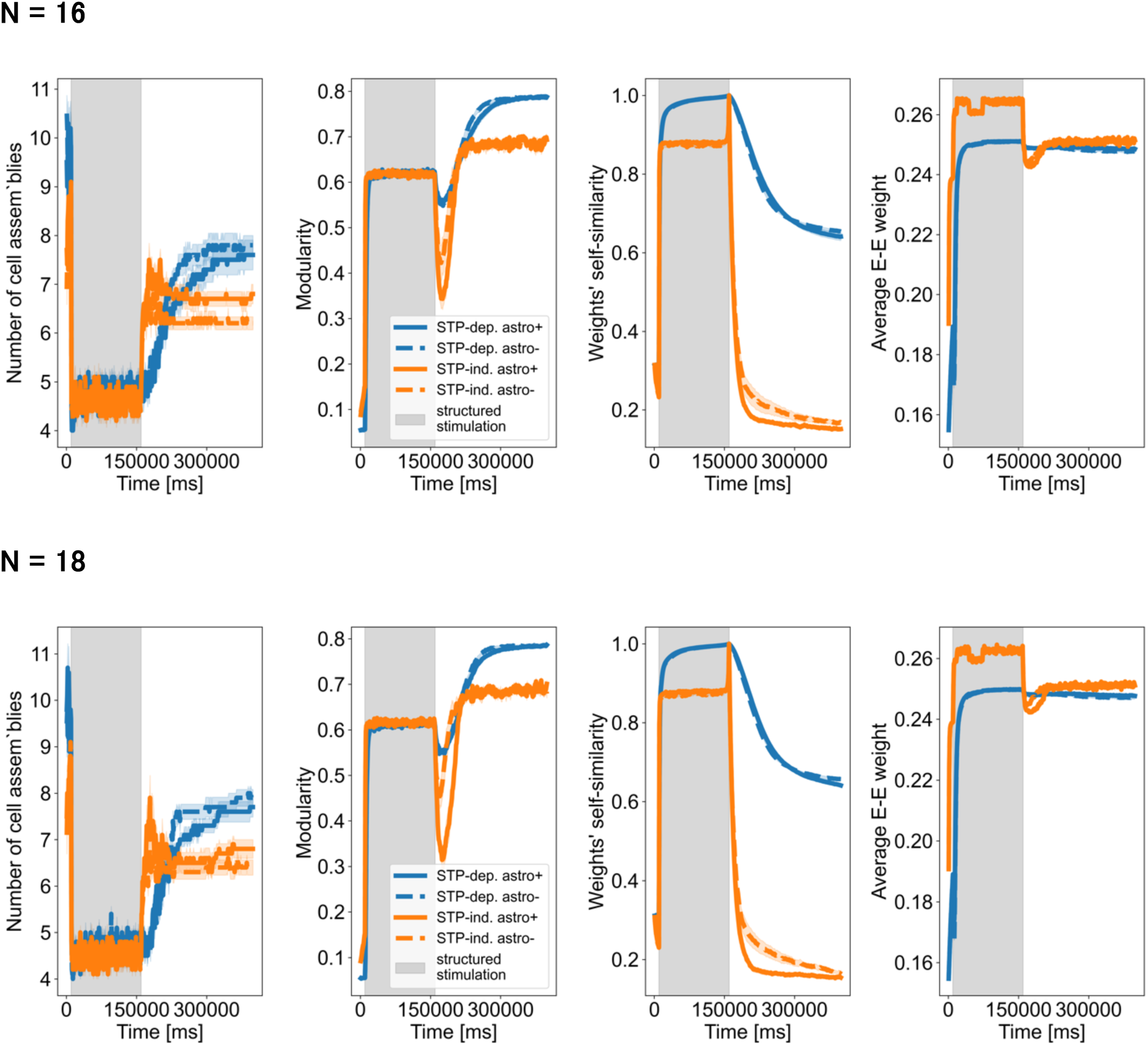
Stability of cell assemblies learned from structured stimulation for *M* ∈ {2, 4, 6, 8, 10, 12, 14, 16, 18}. Attempting to learn *M* ∈ {2,4,14,16,18} results in an unstable cell assembly structure that eventually settles to a configuration with about 7-8 cell assemblies for STP-dependent STDP) and 6-7 cell assemblies for STP-independent STDP.

**Figure S3.**
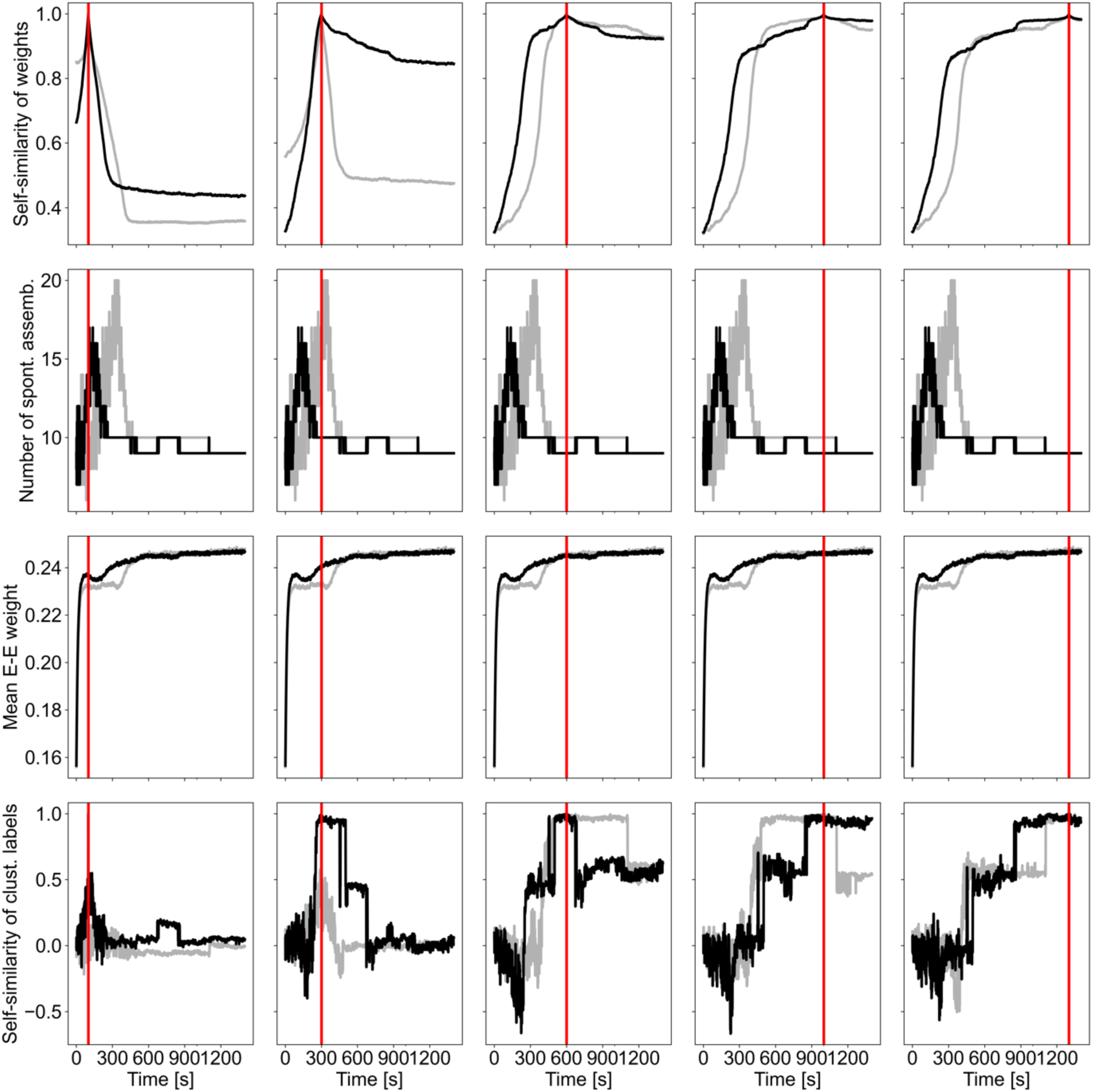
Change of self-similarity over time relative to different reference times. Panels in each row show the evolution of self-similarity of weights and cell assembly labels relative to a reference points in time (100, 300, 600, 1000 and 1300s, shown with red vertical lines).

## Footnotes

1 There sometimes occur short periods in which the number of cell assemblies either increases or decreases by one. These are followed by recovery to the original stable number of cell assemblies.

2 Self-similarity is defined here as the Pearson correlation of the values (of weights or cluster labels) at some reference time with the values recorded at other points of the simulation.

